# Characterization of the pearl millet cultivation environments in India: status and perspectives enabled by expanded data analytics and digital tools

**DOI:** 10.1101/2023.02.18.529051

**Authors:** Vincent Garin, Sunita Choudhary, Tharanya Murugesan, Sivasakthi Kaliamoorthy, Madina Diancumba, Amir Hajjarpoor, Tara Satyavathi, SK Gupta, Jana Kholova

## Abstract

The cultivation of pearl millet in India is experiencing important transformations due to changes in weather, socio-economic trends, and technological progress. In this scope, we propose a new characterization of the pearl millet production environment in India using the latest available data and methodology. For that, we constructed a database incorporating data on various aspects of pearl millet cultivation at the district level from 1998 to 2017. We complemented this analysis using extensive pearl millet agri-system simulations to evaluate crop models’ abilities to reconstruct and analyse the system at an unprecedented scale. We also proposed a new method to infer system parameters from crop model data. Our results show important differences compared to the characterization currently used. The East part of the pearl millet tract (East Rajasthan, Haryana, Uttar Pradesh, and Madhya Pradesh) emerges as the only region where pearl millet cultivation has grown with potential surplus that is likely exported. Important reductions of pearl millet cultivated area in Gujarat, Maharashtra and Karnataka are potentially due to economy-driven transition to other more pro table crops like cotton, maize, or castor bean. The data used also point toward a constant increase of the rain during the growing season which could have major consequences on the future of this crop, with potential positive effects like extra yield but also negative like extra pressure due to more intense and erratic rainfall or transition to more pro table crops requiring more water. Despite difficulties to predict pearl millet yield in rapidly changing environments, the tested crop models reflected reasonably well the pearl millet production system, thus, setting the base for effective system design in future climatic scenarios. Our data and results have been gathered in an open-source interactive online application.

## Introduction

Pearl millet is an important crop cultivated on about 30 million hectares (ha) in more than 30 countries (Jukanti et al., 2016). In India, pearl millet is mainly cultivated under rain-fed conditions during the rainy season (kharif) that represented 92% of the yearly cultivated surface during the period 1998-2017. Due to its capacity to use water efficiently and its heat tolerance, pearl millet is adapted to harsh climatic conditions where other crops fail to produce economic yield (Yadav and Rai, 2013; Yadav et al., 2012). However, despite constant increases across the production area, with an average of 1161 kg/ha over 1998-2017, pearl millet yield remains low compared to other crops in the same regions (e.g. rice: 1967 kg/ha; maize: 1650 kg/ha). Moreover, limited market opportunities and minimum government support are important constraints for farmers to consider pearl millet for cultivation. Thus, pearl millet is mostly used for food and fodder (Nedumaran et al., 2014) and it is under the constant pressure of changes in dietary patterns for crops like wheat or rice (Nagaraj et al., 2013).

The sustainable intensification of complex agronomic systems like pearl millet cultivation in India requires quantitative understanding of production environments which also set the base to effective design of context specific technologies (Hammer et al., 2014; Harrison et al., 2014; Messina et al., 2020). Increasingly, crop simulation modelling (CSM) is used for environmental characterization (envirotyping) and the definition of target population of environments (TPE) (Muchow et al., 1996; Chapman et al., 2000a,b; Chenu et al., 2013). A TPE is composed of environments (TPEs) with more homogeneous biophysical properties. Their identification and use for breeding help to reduce the GxE effect in selection (Braun et al., 1997; Chauhan and Rachaputi, 2014). Therefore, the TPE definition is a prerequisite for several important applications like the optimization of multi-location breeding trials sampling.

The initial characterization of pearl millet cultivation in India dates to 1979 when the National Agricultural Research Project divided the environment into zones A and B. Later, zone A was split into A1 and A parts (Ghosh, 1991; Packwood et al., 1998) (Figure 1A). This zonation mostly relies on the yearly rain pattern and is still considered today as a reference in many pearl millet breeding programs. According to this zonation, pearl millet is cultivated under low precipitations (<400 mm/year) in the north of Rajasthan classified as A1 zone, in the neighbouring regions of south Rajasthan, Haryana, Gujarat, and Uttar Pradesh termed as A zone (>400 mm/year), and central Western India referred to as B zone (>400 mm/year). Except the work by Gupta et al. (2013), no notable revision of the TPE has been proposed recently. That work supported the equivalent to the initial zonation with no difference between the A1 and A zones. However, this analysis was based on yield data of well-managed improved varieties breeding trials. We suggest that a revision of the Indian pearl millet TPE integrating a wider source of information should better capture the complexity of this system (Kholova et al., 2022). This revision is the main objective of the article.

**Figure 1.**
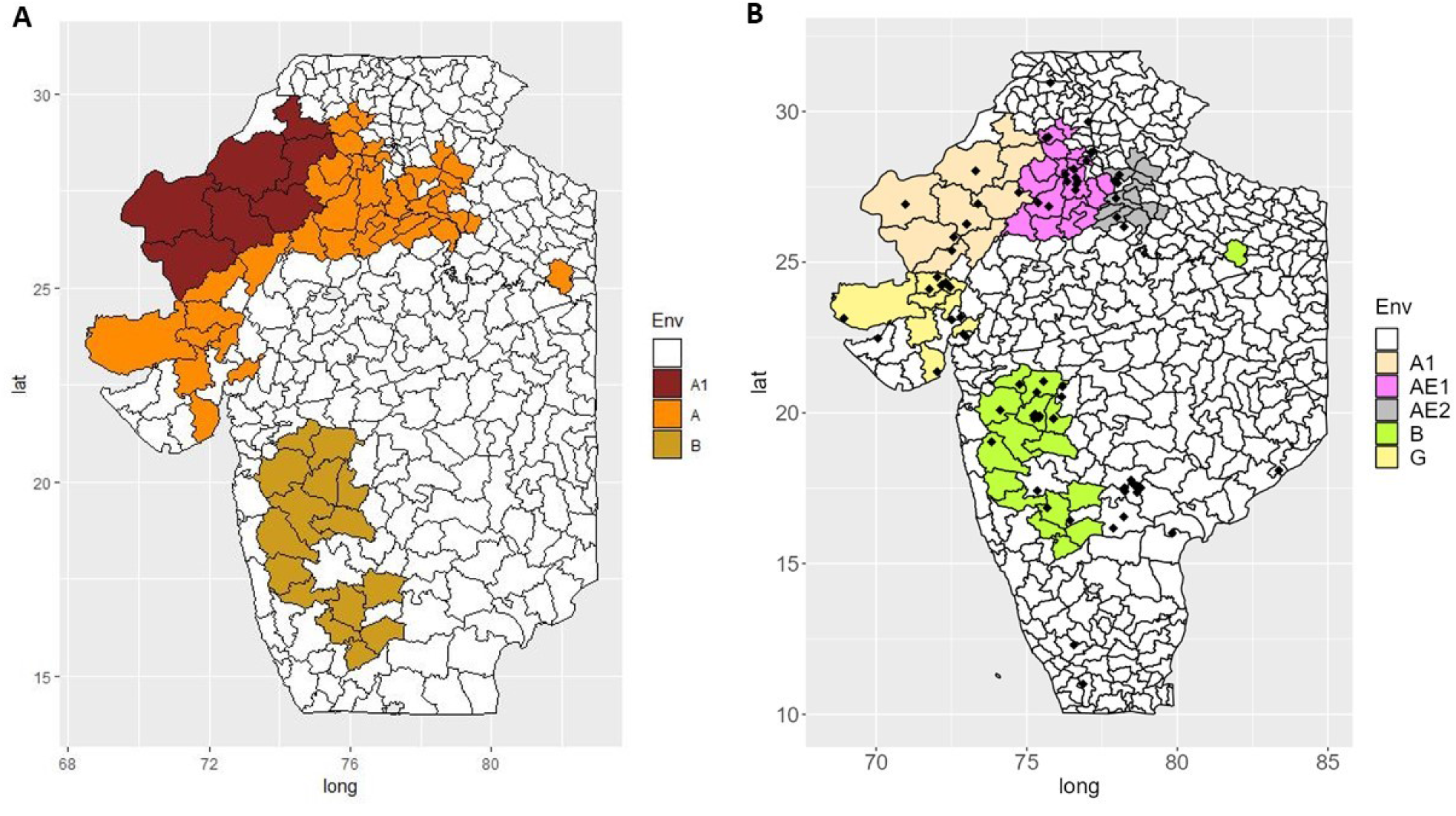
A) Selected 62 districts representing 90 % of the total kharif pearl millet area over the period 1998 - 2017. The districts are coloured according to the reference rain and geography criteria (A1: <400mm/year; A: North >400mm/year; B: South >400mm/year). B) New TPE with AICRP pearl millet testing sites during kharif and summer season 2017

CSM progressively became one of the most important tools for environmental characterization (Kholova et al., 2020, 2022). CSMs are in-silico representations of natural systems captured by a series of equations (Messina et al., 2009). These equations express agronomic traits like yield as a function of environmental inputs (e.g. temperature, soil water potential), management practices (e.g. irrigation, fertilization) and crop characteristics (e.g. phenology, canopy growth, biomass partitioning) (Tardieu et al., 2005). CSM reconstruct the soil-plant-atmosphere continuum of a system across spatio-temporal scales which can be used to identify TPEs with similar biophysical properties forming the TPE.

Recently, increases in computer power opened the possibility to run large scale simulations to extensively explore the properties of agronomic systems (e.g. Ronanki et al. (2022)). The second objective of the article is to present an innovative strategy based on large scale CM simulations to support the revision of the pearl millet TPE. This strategy helped us to gather extra information about the locations where pearl millet is cultivated as well as estimating the influence of different parameters on the yield (sensitivity analysis). This article also allowed us to evaluate a new CSM tool characterized by an improved algorithm to describe pearl millet tillering (Alam et al., 2014, 2017). Since tillering is critical for pearl millet adaptation to drought-prone environment, the evaluation of this updated CSM tool in the new TPE is an important prerequisite to design crops and management strategies for the current and future climates in the target geographies. It will also support pearl millet crop model development, which remains an under-research area (Van Oosterom et al., 2001; Sultan et al., 2013; Singh et al., 2017).

## Material and Methods

Figure 2 is a visualisation of the material and methods workflow. The material and method is divided into two main parts. The first one describe the data used for the TPE determination: a) ICRISAT district level data (DLD); b) Nasapower synthetic weather data (Sparks, 2021); and c) parameters derived from CM simulations (Table 1). The second part describes how the crop model was used to a) infer parameters of the agri-system using large scale simulation, b) use those simulations to evaluate the crop model prediction ability within the identified TPEs and c) determine the relative influence on yield of the tested parameter in the new TPEs.

**Table 1.** Description of the data used given origin (district level data - DLD, Nasapower weather data, crop model inferred parameter) and use in PCA

| Category | Data type | unit | DLD | Weather | Crop model | PCA |
| --- | --- | --- | --- | --- | --- | --- |
| Agronomy<br>(area) | Area | 1000 ha | X |  |  |  |
|  | Area trend | 1000 ha/year | X |  |  |  |
|  | Area share | % | X |  |  | X |
|  | Area share trend | %/year | X |  |  | X |
| Agronomy<br>(production) | Production | 1000 tons | X |  |  |  |
|  | Production increase | 1000 tons/year | X |  |  |  |
|  | Production share | % | X |  |  | X |
|  | Production share trend | %/year | X |  |  | X |
| Agronomy<br>(yield) | Yield | kg/ha | X |  | X | X |
|  | Yield trend | kg/ha/year | X |  |  | X |
| Environment<br>(rain) | Total rain Kharif | mm |  | X |  | X |
|  | Rain variance over year | mm |  | X |  | X |
|  | Rain trend | mm/year |  | X |  | X |
| Environment<br>(temperature) | Min. temperature | °C |  | X |  | X |
|  | Min. temperature trend | °C/year |  | X |  | X |
|  | Max. temperature | °C |  | X |  | X |
|  | Max. temperature trend | °C/year |  | X |  | X |
| Soil | Majority soil type |  | X |  |  | X |
|  | Soil water content | Wat. mm/Soil cm |  |  | X | X |
|  | Soil depth | cm |  |  | X | X |
| Management | Irrigation PM area kh. | % | X |  | X | X |
|  | Irrigation trend | %/year | X |  |  | X |
|  | Fertiliser all crops kh. | kg/ha | X |  | X | X |
|  | Fertiliser trend | kg/ha/year | X |  |  | X |
|  | Improved variety | % cult. area | X |  | X | X |
|  | Sowing date | day |  |  | X | X |
|  | Sowing density | plant/cm<sup>2</sup> |  |  | X | X |
| Price | Price | INR/100 kg | X |  |  | X |
|  | Price trend | INR/100 kg/year | X |  |  | X |

**Figure 2.**
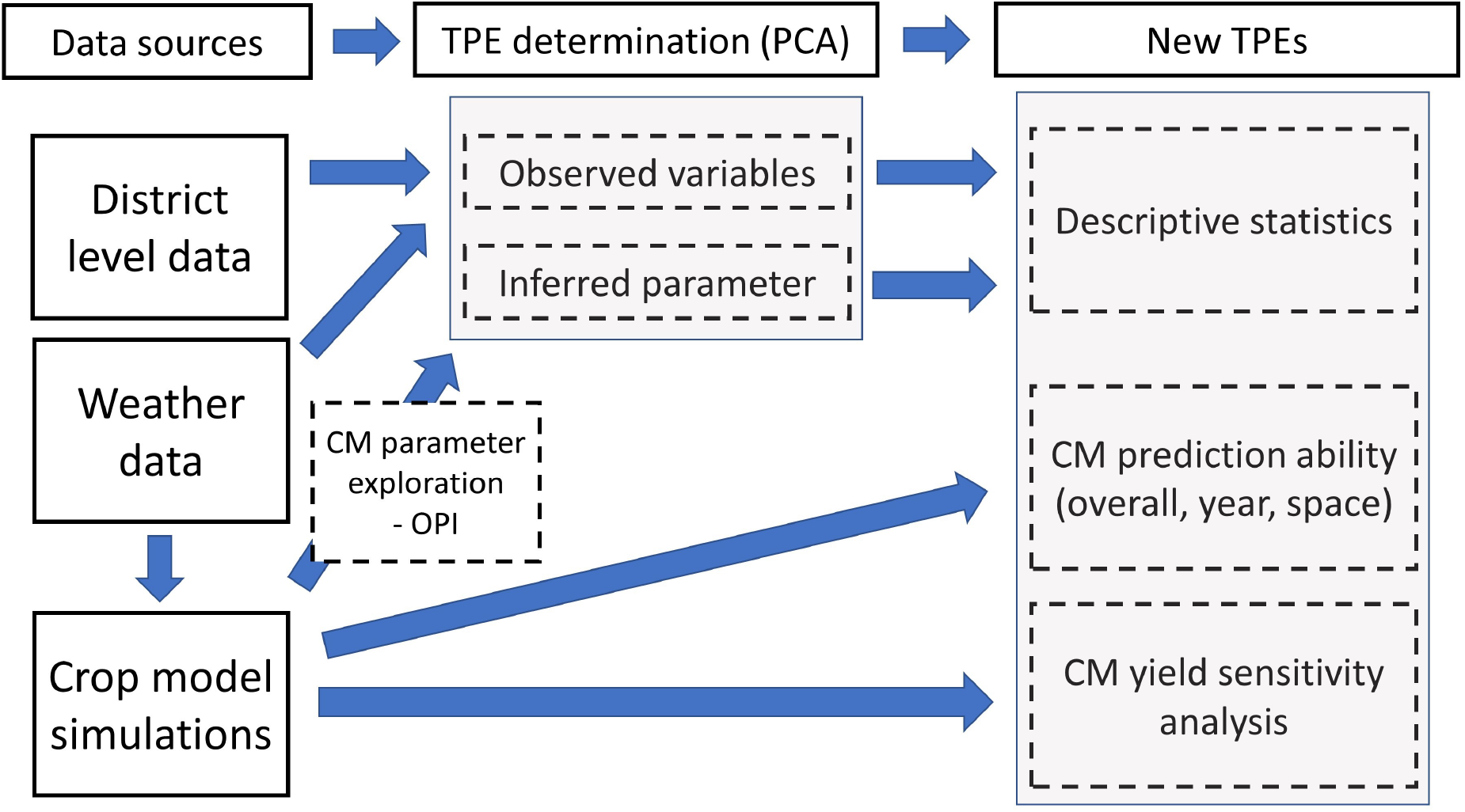
Materials and Methods workflow

### A. Data

#### A.1 District level data (DLD)

The DLD (http://data.icrisat.org/dld/) is an open-source database on Indian agriculture at the district level that contains data originally compiled by the Indian directorate of economics and statistics (https://desagri.gov.in/). From this database, we extracted the most recent 20 years (1998-2017) of available data on pearl millet cultivated area, production, and yield during the rainy season (kharif, Table 1). We identified 62 districts representing 90% of the total cultivated area during the period (Figure 1A). Consequently, the other variables were evaluated for these districts over the selected period. In these 62 districts, we used the area, production, and yield average (1998-2017) and further calculated their increase or decrease over year (i.e. “trend”). For area and production, we also estimated their average share with respect to other crops as the relative proportion and the share trend over 1998-2017. We complemented our dataset with the district majority soil type (Laryea, 1998), the percentage of pearl millet area under irrigation, the amount of applied nitrogen (fertiliser) per hectare for all crops (not available for pearl millet only), and the price at harvest per quintal (Table 1). For irrigation, fertiliser and price we could also estimate the trend.

#### A.2 Nasapower weather data

For weather, we used the synthetic data provided by Nasapower (Sparks, 2021) which has been shown to sufficiently represent the observed weather data (Hajjarpoor et al 2019, Ronanki et al. 2022). The selected data contained daily minimum and maximum temperature, precipitation, and solar radiation that was also used as the CSM input. We used the apsimx R package (Miguez, 2022) to extract the data for a point located at the centre of each district. We calculated the cumulative rain during kharif season (15 June - 15 September) as well as its trend and variance over years (Table 1). We also calculated the temperature trends.

### B. TPE determination

The agronomic and weather data were complemented by parameters inferred from the crop model analysis (Table 1, see next section). Those data were used to perform a principal component analysis (PCA). For area and production, we used the within district proportion (share) rather than the absolute value. We used k-mean clustering with the number of clusters fixed to three, four and five. The districts grouped in the same cluster were considered as more homogeneous than the one in other clusters and were defined as the new TPEs forming the TPE.

### C. Crop model inference strategy

We present a strategy based on extensive simulations to evaluate the CM behaviour and derive information about the system. By generating those outputs, we could a) evaluate the CM ability to reconstruct the pearl millet production at the district level; b) infer system parameters that support the system characterization; and c) determine the main constraints of the system.

#### C.1 Crop model description

We used two versions of the Agricultural Production Systems sIMulator APSIM pearl millet model (Holzworth et al., 2014). The default model was a first attempt to integrate pearl millet in APSIM 7.10 by modelling dynamic tillers as an intercrop of the different axes (Van Oosterom et al., 2001, 2002). The second model extended the default version to reflect more recent biological knowledge and integrated a modifications of canopy developmental dynamics which included the algorithm reproducing the dynamic growth of tillers based on the carbohydrates supply/demand and the tillering propensity of given genotype as proposed by Alam et al. (2014, 2017) (Kim et al., 2010a,b). For a more detailed description of the two CMs see Garin et al. (2023).

#### C.2 Crop model parameters space

A challenge for CM prediction is the determination of the parameters corresponding to the studied system (Wallach et al., 2018). In controlled field experiments, parameters like the soil type or the plant density can be assessed. However, for larger scale predictions as in our case at the district level, the parameters are inferred with less prediction and need to be generalized. Those uncertainties reduce the CM prediction accuracy (Huang et al., 2019). A standard approach is to determine the most likely parameters after literature review and discussion with specialists (Hajjarpoor et al., 2018, 2021). Here we used this approach and called it an “informed strategy”. However, due to the dynamic nature of complex agronomic systems accelerated by climate change, the informed strategy can reach some limits especially in marginalized systems with high heterogeneity. The computation of many simulations defined by various parameter combinations allows the user to not fix the parameters but to determine which parameter values are the most likely by comparing the prediction with real observations. We called this approach optimized parameter inference (OPI).

To evaluate the CM in different parameter configurations, we defined a range of possible values for the main parameters (Table 2). We determined the relevant ranges after DLD analysis, review of the literature and discussions with experts. For the soil parameters, we modified the generic APSIM soil profiles (Koo et al., 2015). We selected three soil textures: sandy, loam, and clay soils, representing the majority soil types (pssament, inceptisol, and vertisol) observed in the pearl millet tract (Supplementary material S1). These soil types are characterized by plant available water capacity (PAWC) of 0.6, 0.9, and 1.3 mm of water per cm of soil, respectively (Burk and Dalgliesh, 2013). We combined each soil texture with three different soil depths of 60, 120 or 180 cm - representing shallow, medium and deep soil profile.

**Table 2.**
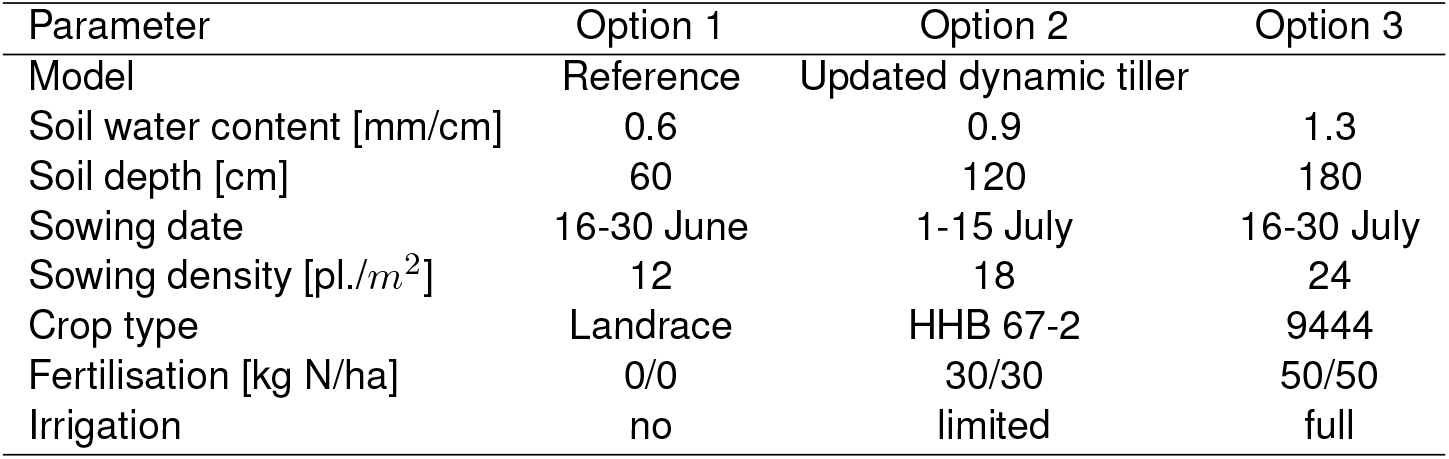
Crop model set-up parameter ranges

| Parameter | Option 1 | Option 2 | Option 3 |
| --- | --- | --- | --- |
| Model | Reference | Updated dynamic tiller |  |
| Soil water content [mm/cm] | 0.6 | 0.9 | 1.3 |
| Soil depth [cm] | 60 | 120 | 180 |
| Sowing date | 16-30 June | 1-15 July | 16-30 July |
| Sowing density [*pl./m<sup>2</sup>*] | 12 | 18 | 24 |
| Crop type | Landrace | HHB 67-2 | 9444 |
| Fertilisation [kg N/ha] | 0/0 | 30/30 | 50/50 |
| Irrigation | no | limited | full |

According to the literature, the generally practiced sowing window is the first two weeks (1-15) of July which, however, is conditioned by the onset of the rains (Rana et al., 2012). Considering these, we defined an early (16-30 June) and a late sowing window (16-30 July). Accordingly, the CM was set to initiate the crop when at least 8.5mm of rain was accumulated within five days and the soil contained a minimum of 25 mm of available water. In terms of plant density, agronomic guidelines advise to sow between 11 to 22 *plants/m*^2^ (Yadav et al., 2012). Therefore, we used 12, 18, and 24 *plants/m*^2^. In terms of crops, we used three crop types: landrace, HHB67-2, and 9444 representing generic plant types used in India. The landrace is representative of local varieties used in the West part of Rajasthan (Van Oosterom et al., 2001). HHB67-2 and 9444 are two hybrids with short and long duration, respectively (Garin et al., 2023). The 9444 crop type was only available in the updated CM.

The DLD about fertilisation in the selected districts indicate that during kharif the average quantity of Nitrogen (N) used ranges from 15.5 kg/ha in the A1 zone to 82.8 in the B zone. To approximate these ranges, we selected: a) no fertilization; b) 30 kg/ha as basal dose + 30 kg/ha at 20 days; and c) 50 kg/ha as basal dose + 50 kg/ha at 20 days. Similarly, for irrigation, the DLD show that pearl millet is mostly rainfed with on average 4.9% of the cultivated surface under irrigation in the A1 zone and 15.7% in the A zone. Therefore, we used: a) no irrigation, b) limited irrigation applied when the fraction of available soil water (ASW) is below 0.25, and full irrigation when ASW fraction is below 0.5.

#### C.3 Crop model optimized parameter inference

CM can be seen as a function 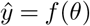 that predicts an output like yield given the parameters *θ* (e.g. the level of irrigation) (Wallach et al., 2018). Generally, CM accuracy depends on its representativity (i.e. the fact that the model integrates all critical biophysical phenomena) and the validity of its functions (i.e. the fact that the bio-physical mechanisms are correctly simulated), the quality of the observations, and the accuracy of the parameters (Huang et al., 2019). Assuming no error in the model and the observations, one can estimate the likelihood of a selected parameter by calculating the CM output 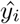 for different values and comparing the output with real observations *y*. The parameter values resulting in the smallest difference 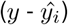 should be the most likely (Supplementary material S2). In our case, for each of the 3645 unique parameter combinations (Table 2) we predicted the yield in each district yearly. We determined the most likely parameter combination as the one with the highest Pearson correlation (*ρ*). Using those results we could obtain continuous values for the parameter inferred by OPI by calculating the weighted average of the parameter value from the 5% best *ρ*. The weights were proportional to *ρ*.

#### C.4 Crop model evaluation and parameter stability over time and space

We evaluated the crop model general prediction ability for each district using the correlation between the yield observed time series (DLD) and the predictions obtained with the inferred optimized parameter (OPI) value combination. We compared the CM prediction ability obtained with the OPI to the one obtained with the informed strategy. For the later, we determined in each original zones (A1, A, and B), the most likely parameter values using available data and literature (Supplementary material S3).

The simulation output also allowed us to evaluate the capacity of selected parameters to predict outcome in pseudo independent data over time and space (parameter stability). For that purpose, we used a strategy similar to cross-validation where we estimated the prediction ability in independent years and locations (Roberts et al., 2017). The year data were split into 1998-2007 and 2008-2017. In the first scenario (time effect), we determined the optimal parameters for a district using the present years and then predicted the yield in future years. In the second scenario, (location effect), we used the parameters determined in one district in the present years to predict the yield of the closest neighbouring district by at least 150 km in the same years. The prediction ability over time and space was evaluated with *ρ*.

#### C.5 Crop model sensitivity analysis

The generated CM output allowed us to perform a sensitivity analysis to evaluate the parameter effect on yield in each newly determined TPE. We used linear regressions of the predicted yield given parameters (e.g. 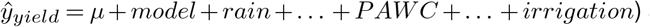 and weather parameters (rain, temperature) to determine the magnitude of their effect and the most influential one in a specific TPE by averaging the results over districts.

## Results

In the results section, we first present the revised TPE and describe each TPE using descriptive statistics. Then, we discuss the CM performances and its behaviour in the newly determined environments.

### D. Characterization of the target population of environments (TPE) and comparison with the current zonation system

Before describing the new TPE (Figure 3), let us mention which variables were the most discriminative to differentiate the districts in the PCA analysis. Table 3 is an exhaustive description of the new TPEs given 40 variables covering surface, production, yield, weather, soil, management, and price. The most important parameters were the weather (rain, temperature) and the soil type and properties (PAWC, depth). Then, the crop area, production, and yield trends were also highly discriminative. Management practices like fertilisation and irrigation, which reflect the level of input, had a more moderate effect. The CM inferred parameters like sowing date characterized by a higher uncertainty compared to the observed data were less influential.

**Table 3.**
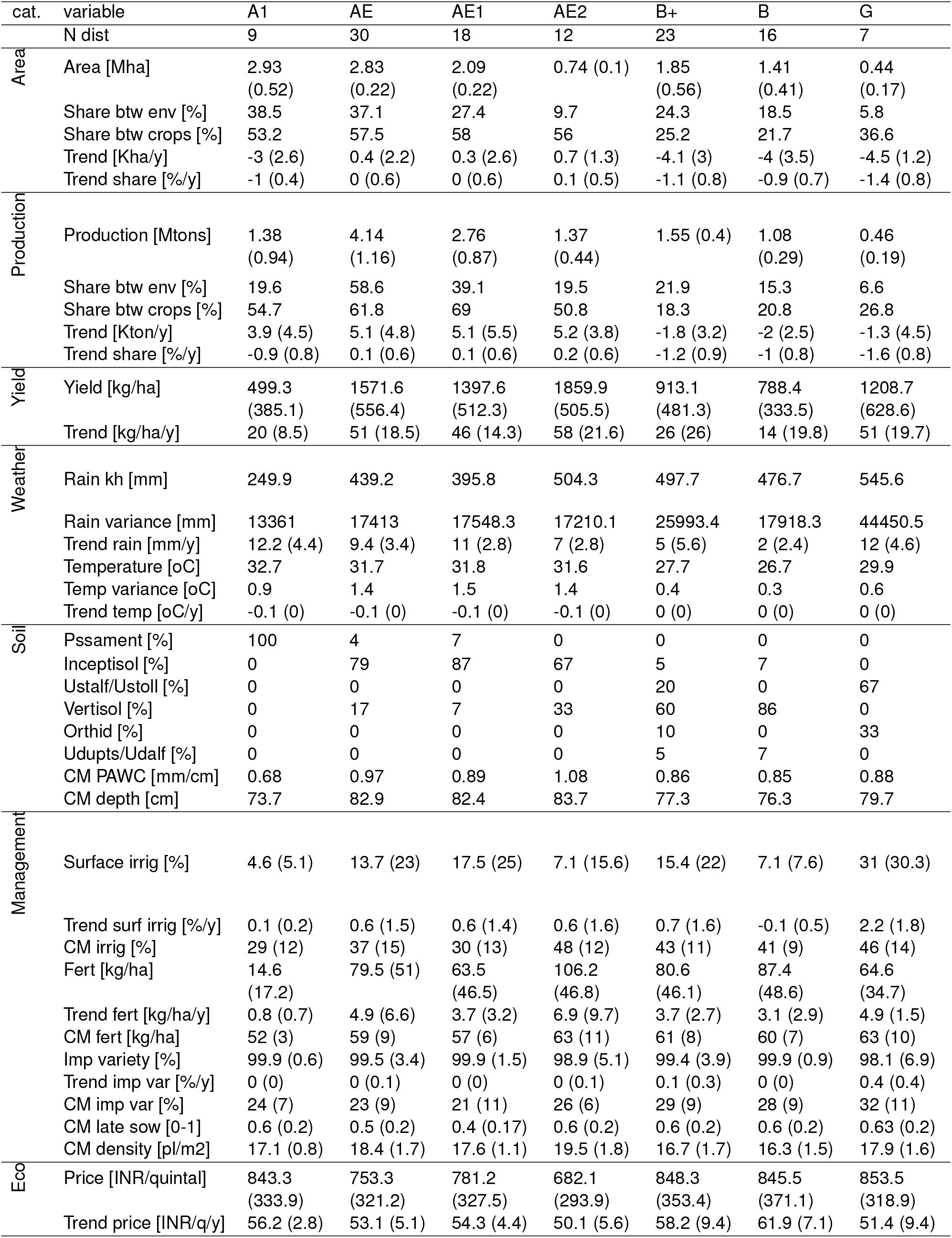
Descriptive statistics for the newly determined TPEs (A1, AE1, AE2, B, G). AE covers (AE1 and AE2) B+ covers (B and G)

**Figure 3.**
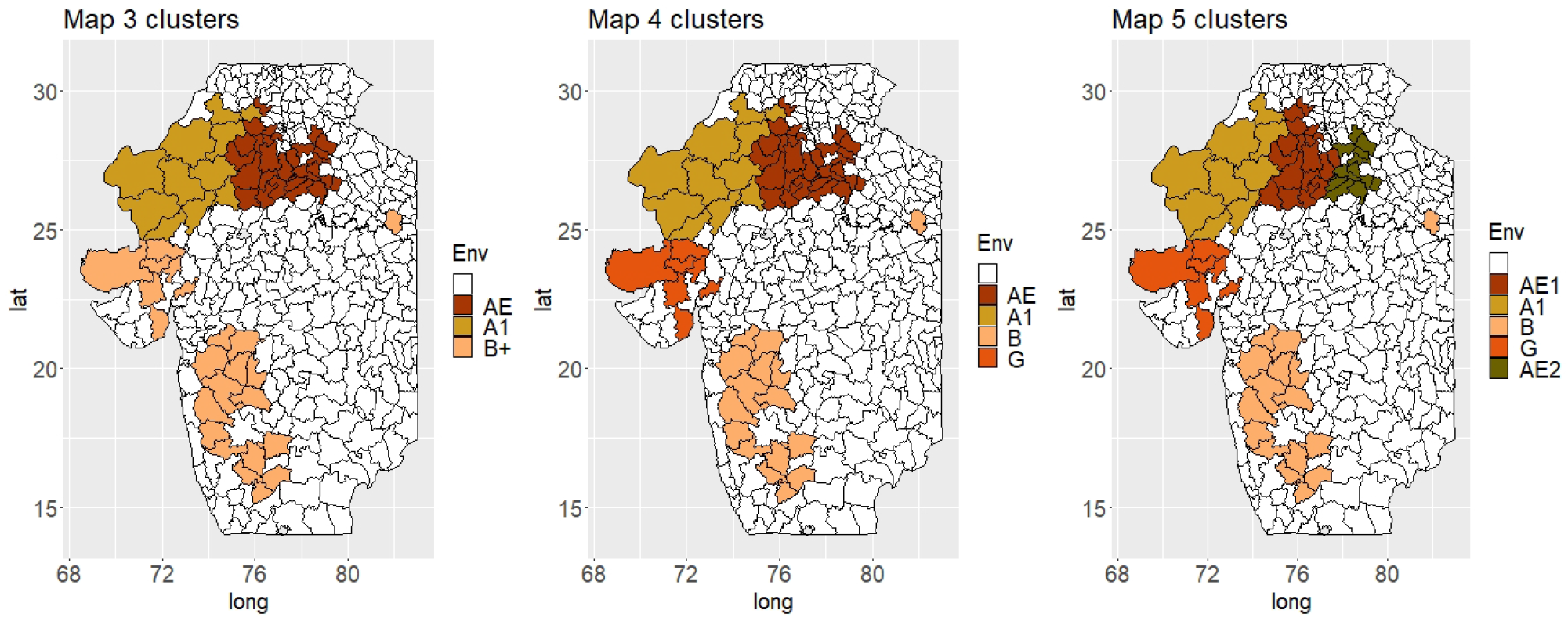
Maps of new zones determined by the PCA clustering with the number of clusters fixed at three, four, or five

The proposed zonation based on TPE characterization showed similarities with the current system for the A1 (North Rajasthan) and B zones that roughly remained the same (Figures 1 and 3). However, the previously defined A zone showed potentially more diversity and can be re-defined in two or three distinct parts (G, AE [AE1, AE2]). Within these, the districts of Gujarat (G) shared more similarities with the B zone and could be seen as a unique entity. Similarly, the East part of the original A zone (AE) could be further split in up to two parts (AE1, AE2).

#### D.1 A-Eastern (AE) TPE

The proposed AE TPE would encompass the North-eastern districts belonging to A zone of original classification (Rajasthan, Haryana, Uttar Pradesh, and Madhya Pradesh). This is the only region where pearl millet cultivation area has increased (Figure 4; average rate of 0.4 Kha per year between 1998 and 2017). The eastern part of AE (AE2) was the only geography where the share pearl millet area compared to other crops has increased (+0.1% per year). The AE TPE was found the most important in terms of production (average 58.6% of the total production) and in terms of area (2.83 Mha on average). The AE districts had reported the highest yields (1572 kg/ha on average) and have experienced an important increase (51 kg/ha/y), especially in AE2 (58 kg/ha/y).

**Figure 4.**
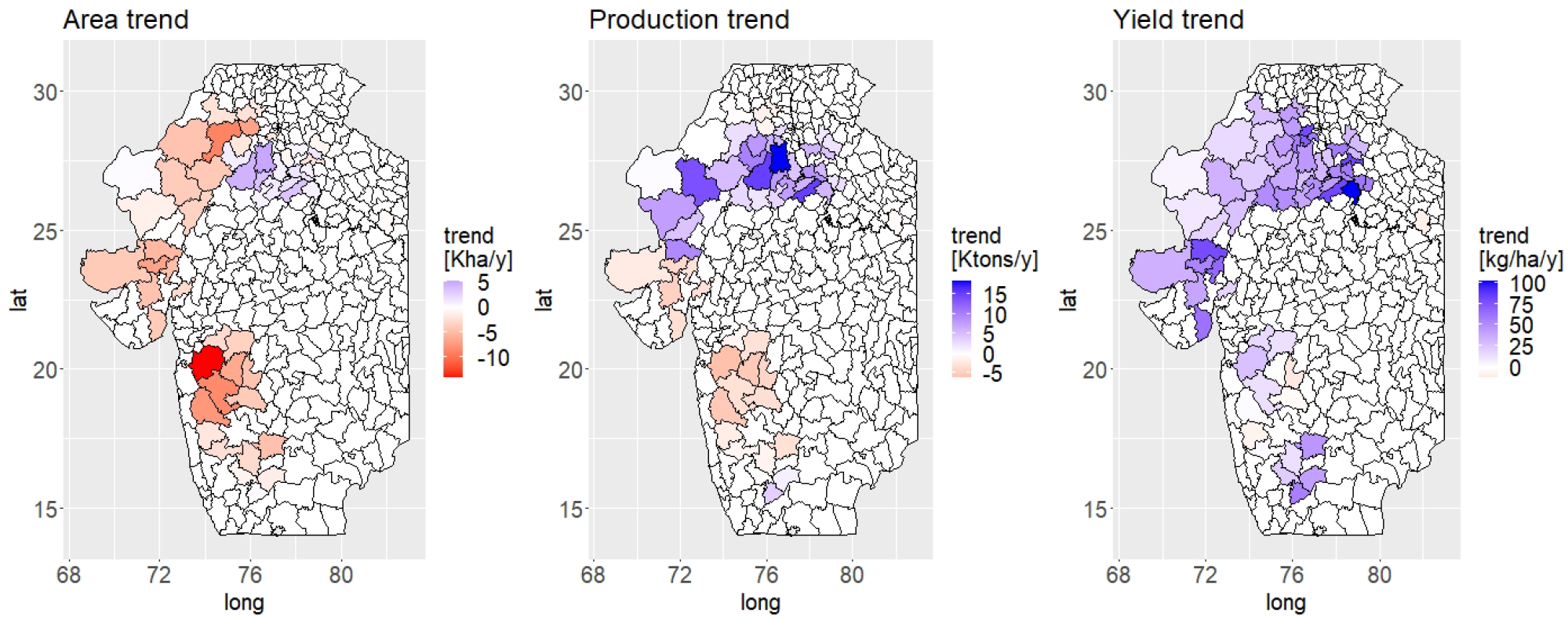
Kharif pearl millet cultivated area, production, and yield trends over 1998-2017

The influence of the favourable weather conditions, soil and management practices can possibly underlie the success and expansion of pearl millet cultivation in AE. As in the other TPEs, according to the weather data, the cumulative rain during kharif has consistently increased over the studied period (average of 504 mm in 1998-2017 with an increase rate of 7 mm per year). AE2 is the TPE with the second highest precipitation. In terms of soil, the inceptisol with higher PWAC (as per OPI) covers most of the AE districts and this further supports the pearl millet production. Finally, in terms of production technology, the AE TPE had a relatively higher proportion of irrigated surface (13.7%) and a relatively high usage of fertiliser with AE2 documented as the largest user of nitrogen (106.2 kg/ha). Part of this success could also be due to a good dissemination of improved varieties in AE (Rao et al., 2018). Finally, the AE TPE is the one with the lowest level or price per 100kg (753 INR), especially in the AE2 TPE (682 INR), which is around 20% less than in A1 (843 INR).

#### D.2 The arid TPE - A1

The TPE characterization recovered most of the original A1 TPE encompassing 11 western districts of Rajasthan. During 1998-2017, the cultivated area decreased on average by three Kha/year while yield moderately increased (20 kg/ha/y), especially in the southern part (e.g. Jodhpur 34 kg/ha/y), which allowed the production to grow (Figure 4). With an average of 2.93 Mha (38.5 % of the total area) this TPE remains the most important in terms of pearl millet cultivated surface. The A1 districts experience the most difficult environmental conditions with the lowest average precipitation (249.9 mm) and a unique type of sandy soil (pssament), which has potentially the lowest PAWC and depth. Those constraints probably explain why the yields remain the lowest in A1 (499 kg/ha) and this goes hand-in-hand with the low recorded irrigation (4.6 % of the area) and fertilisation (14.6 kg/ha) levels.

#### D.3 The transitioning TPE – B

The identified B TPE largely resembles the original B zone, but it has experienced the strongest changes between 1998 and 2017 because the cultivated area and the production have strongly declined (Figure 4). With a yield increase of 14 kg/ha/y, the B TPE had the smallest yield increase. Some districts even experienced a yield decline (e.g. Satara −6.25 kg/ha/y). The main reason explaining this decline might be the competition with alternative crops (Supplementary material S4). Among those, cotton and maize seemed to replace pearl millet in many districts. The decline of pearl millet cultivation can be also linked to a reduction of the resources allocated to this crop (i.e. fertilization, irrigation). For example, the B TPE is the only environment where pearl millet irrigation has declined (−0.1%/y).

#### D.4 The Gujarat TPE – G

The last (G) TPE encompassing seven districts belonging to Gujarat shares some characteristics with the B TPE in terms of decreasing trends and with the AE TPE in terms of increasing yield (+ 51 kg/ha/y) and management practices. Like the B TPE, G has experienced an important decline of cultivated area (−4.5 Kha/y) and production (−1.3 Kton/y) which could be due to surface re-allocation to crops like cotton or castor bean (Supplementary material S5). In terms of weather, the G TPE shows the highest average of in-season rain (545.6 mm) and the strongest rainfall variance. Such a feature could explain a partial re-distribution of the pearl millet production from kharif (rainy season) to the summer (hot season). In terms of production technology, the data show trend different from the B TPE because inputs like irrigation or fertilization have increased over 1998-2017. For example, irrigated surface increased on average by 2.2% per year.

### E.Crop model prediction ability

Table 4 summarizes the CM capacity to reflect pearl millet production in identified TPEs, over time, and over space. The overall CM yield prediction ability (*ρ*) was the largest in AE1 (0.54-0.78) and A1 (0.51-0.76) and the lowest in AE2 (0.18-0.61) and B (0.03-0.52). The yield prediction over years was also the largest in AE1 (0.32-0.46) and A1 (0.28-0.36) compared to AE2 (−0.06 - −0.02) and B (−0.14-0.09). This is potentially due to stronger changes in pearl millet production over years in B, AE2, and G compared to A1 and AE1 (see above). Concerning the location effect, the *ρ* values were comparatively lower in the B TPE, which suggests a relative environmental uniformity within the TPEs, except B. Such a stronger variability of the system parameter over time compared to space was expected.

**Table 4.** Crop model prediction ability (Pearson correlation - *ρ*) over TPEs in the different tested scenario (overall, year, location)

|  | meth | A1 | AE1 | AE2 | B | G |
| --- | --- | --- | --- | --- | --- | --- |
| Overall | Informed | 0.51 | 0.54 | 0.18 | 0.03 | 0.39 |
|  | OPI | 0.76 | 0.78 | 0.61 | 0.52 | 0.67 |
| Year effect | Informed | 0.36 | 0.46 | -0.06 | -0.14 | 0.28 |
|  | OPI | 0.28 | 0.32 | -0.02 | 0.09 | 0.08 |
| Location effect | Informed | 0.54 | 0.61 | 0.49 | 0.05 | 0.72 |
|  | OPI | 0.53 | 0.45 | 0.29 | 0.23 | 0.56 |

The overall prediction ability obtained with the OPI method (0.52-0.78) was greater than the one obtained with the informed strategy (0.03-0.54). However, in the evaluation of year and location effects, the informed strategy gave more stable results than the OPI except in the B TPE where the informed strategy failed. This result illustrates the advantage of the OPI strategy for locations where information is incomplete, or the conditions are rapidly changing like the B TPE. However, the OPI strategy tends to overfit the data which limits its capacity of prediction in time and space while the informed strategy seems to offer more stable results.

### F. Ranges of crop model parameters and weather input influence on yield

In Table 5, we synthesized the influence of each studied input CM parameter range (Table 2) on simulated yield in the different TPEs. Overall, the soil water content (PAWC, 0.6 - 1.3 mm/cm soil) was the parameter with the strongest consistent effect (*R*^2^ *∈* [0.06*−* 0.1]). Soils with low water (0.6) content had significantly reduced yield (600-730 kg/ha) compared to high water content (1.3). The crop types also had an overall strong effect on yield. The improved long duration crop type (9444) had generally a positive effect on yield (617-938 kg/ha) except in the A1 TPE where the effect was less important (342 kg/ha). In that TPE, the effect of tested irrigation regimes was the strongest (*R*^2^ = 0.17) with an advantage of 4.75 kg/ha/mm. Irrigation effect was less important in the other TPEs. Compared to soil PAWC, crop type, and irrigation other ranges of tested management practices like fertilisation, sowing date, or sowing density had less influence on yield.

**Table 5.** Crop model parameter influence on grain yield in the new TPEs as the estimated linear regression coefficient (B) and the R squared statistics over all simulations

| var. | A1 |  | AE1 |  | AE2 |  | B |  | G |  |
| --- | --- | --- | --- | --- | --- | --- | --- | --- | --- | --- |
| | B | $R^2$ | B | $R^2$ | B | $R^2$ | B | $R^2$ | B | $R^2$ |
| Cum rain | 3.13 | 0.03 | 1.15 | 0.01 | -0.30 | 0.00 | -0.41 | 0.02 | -1.14 | 0.02 |
| MaxT | -274.66 | 0.03 | -326.16 | 0.05 | -205.19 | 0.02 | -250.04 | 0.01 | -410.39 | 0.07 |
| MinT | 244.54 | 0.02 | 194.69 | 0.01 | 101.24 | 0.00 | 489.15 | 0.02 | 511.27 | 0.04 |
| Model:new | 0.00 | 0.03 | 0.00 | 0.06 | 0.00 | 0.08 | 0.00 | 0.18 | 0.00 | 0.13 |
| Model:old | -310.94 |  | -515.09 |  | -656.03 |  | -1050.82 |  | -810.35 |  |
| PAWC:1.3 (clay) | 0.00 | 0.10 | 0.00 | 0.10 | 0.00 | 0.09 | 0.00 | 0.06 | 0.00 | 0.10 |
| PAWC:0.9 (loam) | -317.95 |  | -212.49 |  | -54.22 |  | -40.73 |  | -163.03 |  |
| PAWC:0.6 (sand) | -719.26 |  | -712.41 |  | -633.46 |  | -600.34 |  | -732.02 |  |
| SoilDepth:180 | 0.00 | 0.03 | 0.00 | 0.03 | 0.00 | 0.02 | 0.00 | 0.04 | 0.00 | 0.02 |
| SoilDepth:120 | -10.75 |  | -4.99 |  | -3.05 |  | -5.17 |  | -3.64 |  |
| SoilDepth:60 | -348.39 |  | -324.55 |  | -297.91 |  | -464.91 |  | -325.53 |  |
| Sowing:early | 0.00 | 0.00 | 0.00 | 0.01 | 0.00 | 0.01 | 0.00 | 0.00 | 0.00 | 0.00 |
| Sowing:average | -33.06 |  | -104.43 |  | -141.86 |  | 13.67 |  | -107.31 |  |
| Sowing:late | -126.58 |  | -293.17 |  | -422.51 |  | 65.67 |  | -179.60 |  |
| Density:12 | 0.00 | 0.00 | 0.00 | 0.00 | 0.00 | 0.00 | 0.00 | 0.00 | 0.00 | 0.00 |
| Density:18 | 2.89 |  | 22.94 |  | 27.49 |  | 59.15 |  | 37.77 |  |
| Density:24 | -21.50 |  | 4.50 |  | 10.95 |  | 63.99 |  | 33.25 |  |
| Variety:Landrace | 0.00 | 0.02 | 0.00 | 0.06 | 0.00 | 0.11 | 0.00 | 0.11 | 0.00 | 0.08 |
| Variety:HHB67 | 4.25 |  | 7.21 |  | 81.74 |  | -20.67 |  | -50.24 |  |
| Variety:9444 | 342.73 |  | 616.67 |  | 928.79 |  | 938.29 |  | 727.95 |  |
| Fertilizer:0 | 0.00 | 0.00 | 0.00 | 0.00 | 0.00 | 0.01 | 0.00 | 0.01 | 0.00 | 0.01 |
| Fertilizer:60 | 26.86 |  | 68.38 |  | 129.68 |  | 134.92 |  | 113.70 |  |
| Fertilizer:100 | 41.91 |  | 117.21 |  | 236.16 |  | 241.59 |  | 208.92 |  |
| Irrigation [mm] | 4.75 | 0.17 | 4.14 | 0.05 | 3.08 | 0.01 | 5.35 | 0.02 | 3.79 | 0.02 |

Concerning the effect of the weather parameters, we noticed that extra maximum temperature consistently reduced yield while extra minimum temperatures increased it. Interestingly, an extra rainfall had a positive effect on simulated yield in A1 and AE1 but a negative one in the G, B and AE2 TPEs. Finally, we notice a strong influence of the model used with *R*^2^ values between 0.03 and 0.18. The updated version of the model tended to overestimate the yield. However, the updated model was better to predict the yield trend in the A1 and AE1 TPEs (Supplementary material S6).

### G. Detailed results about OPI strategy

We found that OPI could predict the observed data but, at the same time, it had a tendency to overfit them, which resulted in lower prediction of independent year and locations. Another evaluation for the OPI strategy was to compare the parameters values inferred by OPI with the actual observations. In Figure 5, we compared the irrigation and fertilization OPI parameters to the observed percentage of irrigated surface and level of N fertilisation (DLD database, all crops). In both cases, we noticed that the observed and OPI-inferred parameters had a similar spatial distribution across the investigated districts. Irrigation was high in the eastern and Gujarat districts. However, in the B TPE the OPI overestimated irrigation. Concerning fertilisation, the recorded usage of nitrogen was consistently higher in the eastern districts of as well as the B TPE in both observations and OPI parameters. The moderate correlation between OPI parameter and observation of 0.2 and 0.22 for irrigation and fertilisation, respectively showed that the OPI method is only able to approximate the general tendency but fail to predict more extreme observations.

**Figure 5.**
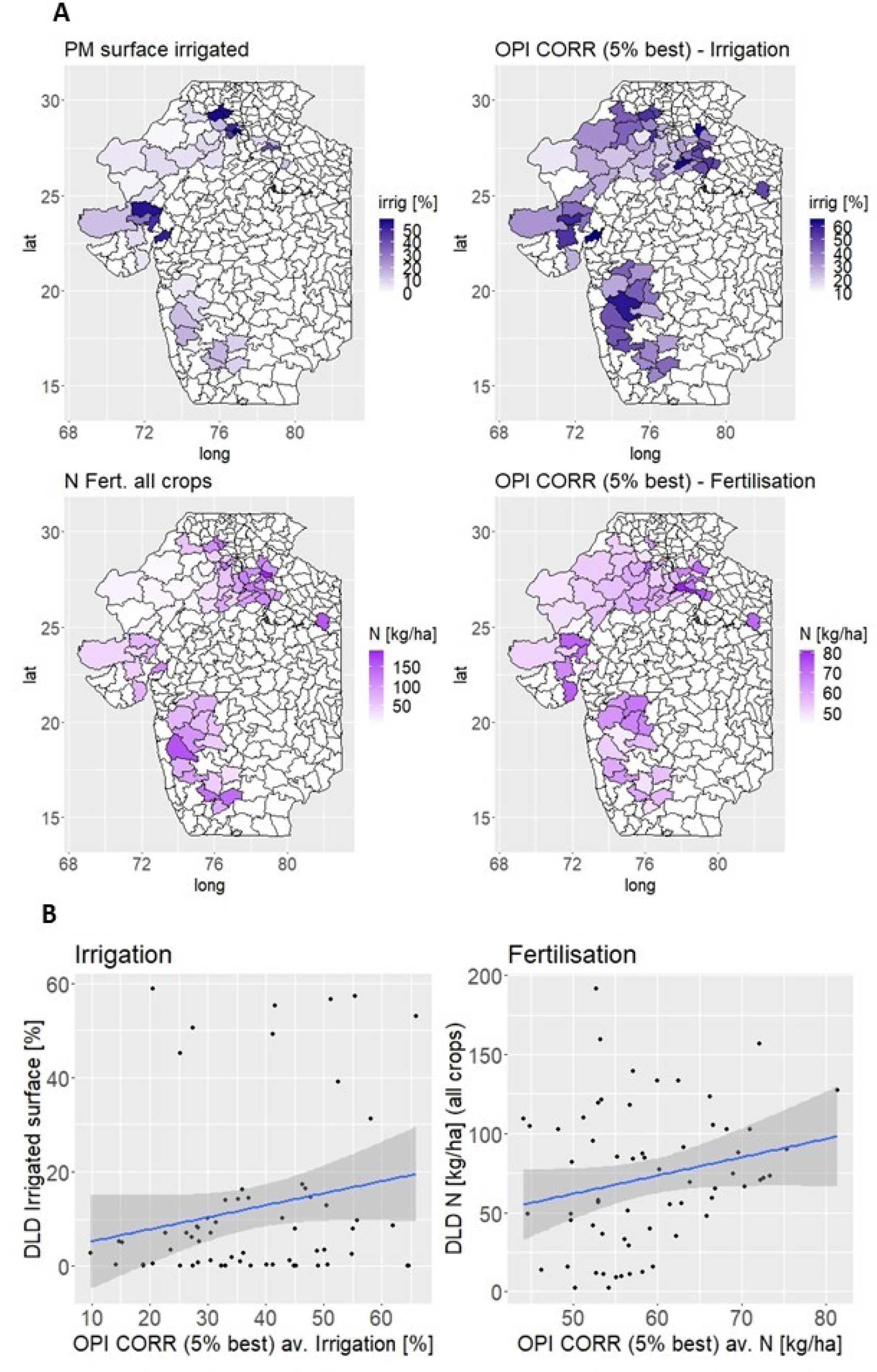
A. Comparison of the OPI parameter values for irrigation and fertilisation with the average percentage of pearl millet surface under irrigation and the average amount of N fertilisation for all crops during kharif according to DLD (1998-2017) B. Scatter plots of the OPI and observed district values for irrigation and fertilisation

The evaluation of the OPI inferred CSM parameters reflecting the soil properties and other management practices (e.g. sowing date) was more difficult because we could not directly compare them to real observations. Nevertheless, we could observe that the inferred soil PAWC and depth followed the trend expected in the observed soil properties (Supplementary material S7). For the other management parameters (sowing date, plant density) with lower influence on yield according to the CM, it was more difficult to identify a meaningful pattern (supplementary material S8).

### H. Online application for results visualisation

We compiled the article data and results in an (online) R shiny application (https://github.com/vincentgarin/PMapp and https://agrvis.shinyapps.io/PMapp/) allowing the user to access them in an interactive way.

## Discussion

The main objective of this article was to characterize the Indian pearl millet production regions using novel data sources and state-of-the-art methodologies. This analysis was originally requested by the Indian national and international research institutions to provide deeper understanding and quantitative base for the optimization of pearl millet production systems. The revised TPE is still largely based on influential biophysical factors like rain pattern, temperature, and the soil characteristics but provides several novel insights, particularly hinting to the changing climate and socio-economic drivers, which should be further investigated to develop successful pearl millet system improvement strategies. In the following sections, we will first discuss the implications of important causes of changes like the possible rain pattern increase and the socio-economic aspects potentially driving some of the TPE features. Then, we will discuss pro- and cons-for the use of CM for pearl millet envirotyping.

### I. Kharif rain increase and transition of pearl millet cultivation to summer season

In all TPEs, the weather data suggests that the amount of precipitation during kharif has increased during the last two decades. The hypothesis of increasing kharif rain is supported by other sources and is expected to continue (Katzenberger et al., 2021). Such a change could strongly influence the future strategy for pearl millet systems improvement and should be closely monitored. We suggest that a part of the yield increase observed over the studied period might be due to those extra precipitations.

In other case however, the potential extra kharif precipitation might directly or indirectly reduce the area allocated to pearl millet cultivation. A first hypothesis assumes a direct negative effect of rain due to an increase of the episodes of high intensity. The extra variability of rain particularly strong in the G TPE could result in short and intense rain episode causing problem of water logging, stem mechanical break and/or grain mold with strong impact on the crop. Another hypothesis assumes an indirect effect. Indeed, we can also imagine that the extra available water during the traditional pearl millet growing season motivate the farmers to switch to crops that require more water like cotton or maize. Such a hypothesis is supported by the important increase area allocated to those crops in the B and G TPEs which could explain the strong decline of pearl millet in those TPEs (see next section).

This direct and indirect effect on rain could explain the transfer of the pearl millet cultivation from kharif to summer season observed in TPE G and to a lesser extent in AE2 (Supplementary material S10). This hypothesis was confirmed by several experts and the DLD data. Indeed, in the G TPE, the proportion of area cultivated in summer has progressively increased to reach more than 50% (Supplementary material S9). However, according to the DLD data, it does not represent a net area transfer between seasons but a larger propensity to cultivate pearl millet during summer because even in summer pearl millet area stagnated or even decrease. In TPE AE2 too, we could observe an increase of the area cultivated in summer with up to 20 % of the yearly surface around 2009. But after, this proportion decreased. Those observations need to be confirmed over longer periods.

### J. The essential role of socio-economic factors in pearl millet cultivation

The average yield increase of 37 kg/ha/year over all districts similar to Yadav et al. (2021) findings hides important differences between TPEs (A1: 20 kg/ha/year, AE2: 58 kg/ha/year) that let assume some differences in the assimilation of new varieties and management practices. Generally, except in the G TPE, the yield increases were correlated with area trends. In TPE B and G, the decline in the pearl millet cultivated area was also strongly correlated with the increased cultivation of high value crops like cotton, maize or castor bean having a more structured value chain and a higher profitability (Blaise and Kranthi, 2019; Hellin and Erenstein, 2009). Therefore, the overall system dynamic seems to be closely related to socio-economic factors. If this is true, efforts to improve pearl millet production could be in vain if farmers continue to replace pearl millet by higher value crops. The necessity to strengthen the pearl millet value chain by increased profitability should therefore be a priority. However, such task seems complicated because government policies as well as trends like urbanization and the changes in dietary patterns are very difficult to influence (Basavaraj et al., 2010).

Another example of the importance of socio-economic factors was the increase of pearl millet cultivation area in the AE2 TPE while the price per quintal was the lowest. According to regional experts substantial parts of AE region production (especially AE2) might be exported to Rajasthan where pearl millet is a staple food and market is more important while regional production possibly not sufficient (OP Yadav personal communication). The increased prices occurring in A1 (around 20 %) might motivate the export between regions. This would represent an example of commercially viable pearl millet value-chain, but the hypothesis should be confirmed by further investigations. Generally, our study emphasizes the existence of different socio-economic drivers for pearl millet cultivation in TPE like A1 that seems more oriented toward subsistence and AE2 which look more market oriented.

### K. The use of crop model for pearl millet envirotyping and its limitations

A second objective of this article was to test the ability of CM to reflect the dynamics of the pearl millet production systems in India and incorporate some of these components into the TPE analysis, which can be a basis for future system optimization (e.g. Ronanki et al. (2022)). Here, we shall emphasize that the APSIM CM do not simulate important biotic stresses like pest and disease attack. For example, it is documented that an important share of pearl millet has suffered from increasing blast attack (Singh et al., 2018). The combined effect of those stresses with the generalisation assumption to represent whole districts can explain an important part of the difference between observation and CM predictions. Despite those, in the A1 AE1 and G TPEs which represent the large majority of area (66%) and production (59%), we could obtain prediction abilities between 0.15 and 0.61 (*R*^2^ squared correlation for comparison) that were comparable to the values about regional prediction from other studies Chen et al. (2018) (0.26-0.42) de Wit et al. (2012) (0.24-0.65).

Nevertheless, in regions that potentially experience rapid changes (B, AE2, and G), the CM was less effective at capturing the production fluctuations. In those TPEs, the prediction ability over the years was strongly reduced, which is likely due to a rapid and large variability in the crop and crop management practices between the seasons. The socio-economic drivers and the change in the rain pattern (see above) could be major drivers of those changes. Therefore, we shall emphasize the interest of combining a large set of data allowing us to understand socio-economic trends that seem to have a strong effect on the ability to model the system using CSM. Reduced prediction ability in AE2, B and G could also result from choice of the testing locations to develop the models (Telangana and Rajasthan). The collection of extra data on crop type in AE2, G and B testing sites during future developments would help to improve the CM prediction ability there.

The evaluation of an updated version of the APSIM model for dynamic tiller simulation was promising. We could show that despite the general overestimation of the average yield, this new model offered more sensitivity to model yield trends (+0.18-0.19 correlation) in the A1 and AE1 (core of the pearl millet tract, 66% area, 59% production). This model could, therefore, be used for optimization of the Indian pearl millet system. The CM sensitivity analysis emphasized the importance of the soil parameters for yield. It supports the statement from Carberry et al. (2009) that this component should get extra attention. Finally, the large effect of the crop type, especially in AE2, B and G suggest to further dissect what might be the optimal genotype properties to enhance the crop production in those TPEs. Extra work on the crop model could also help to clarify the observed negative effect of extra precipitation in AE2, G and B TPEs.

One of the novelties presented in this article was the use of extended simulations to infer parameters that maximized the observed yield prediction (OPI). We showed that this strategy could help to infer parameters with large influence on yield like soil water content, irrigation, or fertilisation, but its capacity to infer less influential parameters like the sowing date was weak. Generally, the OPI method tends to determine parameter values that overfit the observed data. Nevertheless, the OPI method could be useful in situations with low levels of information to complement parameter determination using the literature or the knowledge from specialists.

### L. Summary of the salient findings

In this final section, we summarize the main characteristics for each TPE and provide our insight.

#### A1 (2.9 Mha, 1.4 Mton, 499 kg/ha)

This TPE is characterized by harsh cultivation conditions due to low precipitation, high temperature and potentially poor soils (sandy soils, low water content). This scarcity largely explains the reduced number of alternatives to pearl millet cultivation in A1, which is a reason for its stability there. The environmental conditions can also explain the reduced effect of improved varieties. Potential reduced benefits could explain the low level of input (irrigation, fertilisation) observed in the A1 TPE. Different breeding strategies could be possible. Either developing the varieties adapted to low input conditions or assessing the potential for intensified production practices and high input cultivars in parallel with efforts to increase farmers capacity to realize these in practice. Since our data revealed that a proportion of pearl millet might be imported from AE (further investigation is needed to confirm this hypothesis), increasing the production to meet this demand could be a driver for this process. We also noticed that the A1 TPE is possibly under-represented within the national testing system as it contains only eight sites out of 76 multi-location AICRP testing sites (Figure 1B). Resources from testing sites falling in none of the TPEs (17/76), from the B TPE (17/76) or the G TPE (14/76), which are proportionally over-represented could be re-allocated to A1.

A1 districts (Rajasthan): Barmer, Bikaner, Churu, Hanumangarh, Jaisalme, Jalore, Jodhpur, Nagaur, Pali **AE1 (2.1 Mha, 2.8 Mton, 1397 kg/ha)**: This TPE is by far the area where most of the pearl millet was produced between 1998 and 2017 (around 39% of the total production). The production increase is due to a moderate growth of the cultivated area coupled with high yield increase that appeared to be associated with the rain increase and technological investments (irrigation, fertilization, and possible adoption of new varieties suiting intensified cultivation). We recommend reinforcing attention on this TPE for example by maintaining or increasing the number of AICRP testing sites, especially in the southeast part of the TPE (Figure 1B).

AE1 districts (Rajasthan): Ajmer, Alwar, Bharatpur, Dausa, Jaipur, Jhunjhunu, Karauli, Sawai Madhopur, Sikar, Tonk

AE1 districts (Haryana): Bhiwani, Gurgaon, Hisar, Jhajjar, Jind, Mahendragarh, Rewari

AE1 districts (Uttar Pradesh): Mathura

#### AE2 (0.7 Mha, 1.37 Mton, 1859 kg/ha)

AE2 is the only TPE where the pearl millet relative area has increased between 1998 and 2017. This success could be linked to moderate rain increase and soil type (higher water content and depth) suiting pearl millet cultivation. It is a region where pearl millet production benefited from technological investment (fertilisation, irrigation, possibly adoption of new hybrid cultivars) which made pearl millet competitive with respect to other crops (e.g. pearl millet: 1801 kg/ha; maize: 1896 kg/ha). The pearl millet prices obtained in the TPE are the lowest which support the hypothesis that in AE2 pearl millet might be produced for export into the A1 TPE where pearl millet is mostly cultivated for subsistence. With six testing sites out of 76, this TPE is the least represented among the AICRP testing sites (Figure 1B). Given the agronomic success observed there, it would be strategic to increase the coverage there to understand better the reasons of this success and the way the economic drivers are set there.

AE2 districts (Uttar Pradesh): Agra, Aligarh, Auraiya, Budaun, Etah, Etawah, Firozabad, Kasganj, Sambhal AE2 districts (Madhya Pradesh): Bhind, Morena

AE2 districts (Rajasthan): Dholpur

#### B (1.4 Mha, 1.1 Mton, 788 kg/ha)

The B TPE is characterized by the largest decline of pearl millet cultivation area. The abundance of high value crop alternatives connected to potentially more developed value chains (e.g. cotton and maize) might be the strongest driver leading to pearl millet area reduction. This declining interest in pearl millet cultivation is reflected in a low level of technological investment and resulted in very limited yield increase. We might safely assume that any investment to improve pearl millet production in B might have minimal return unless the crop becomes more profitable compared to alternatives. Therefore, we recommend investigating the reasons for abandoning pearl millet cultivation there.

B districts (Maharashtra): Ahmednagar, Aurangabad, Beed, Dhule, Jalgaon, Jalna, Nashik, Pune, Sangli, Satara

B districts (Karnataka): Bagalkot, Bijapur, Gulbarga, Koppal, Raichur

B districts (Uttar Pradesh): Allahabad

#### G (0.4 Mha, 0.5 Mton, 1208 kg/ha)

In Gujarat, like in the B TPE, the pearl millet production sharply decreased probably due to the competition of more profitable crops like cotton or castor beans. The data suggest that part of the G TPE traditional kharif production has been transferred to summer which, however, does not compensate for the overall decline of the pearl millet production across all the seasons. This transfer is potentially due to increase of rain volume and variability during kharif. Despite relatively high investment in technology (irrigation, fertilization) and high yield, the reduction in the pearl millet cultivated area was not compensated by increased yields, which resulted in overall production decline. Perhaps, this TPE can be used as a testing site to understand the influence of rain pattern on pearl millet cultivation, especially the potential benefits related to transfer between the seasons.

G districts (Gujarat): Banas Kantha, Bhavnagar, Kachchh, Kheda, Mahesana, Patan

## M. Acknowledgments

We thank Dr Greg McLean Dr Erik van Oosterom and Professor Graeme Hammer for their help with the implementation of the new pearl millet model. We also thank Dr P Janila, Dr Ashok Kumar from ICRISAT, Dr OP Yadav and Dr Satyavathi from ICAR, Dr Mahala from SeedWorks international private limited, Dr Verma from Kaveri seed company limited and Dr Sanjana from IIMR for the discussion about the article content.

## P. Authors contributions

TM, SK, MD, JK, SC: Data collection for crop model parametrization VG, JK, SC: Database construction and data analysis VG, JK, SC, SKG, TS: Manuscript writing, revision and editing JK, SC, SKG: Project coordination

## S1: Map of the majority soil type in the pearl millet tract millet tract

**Figure 1:**
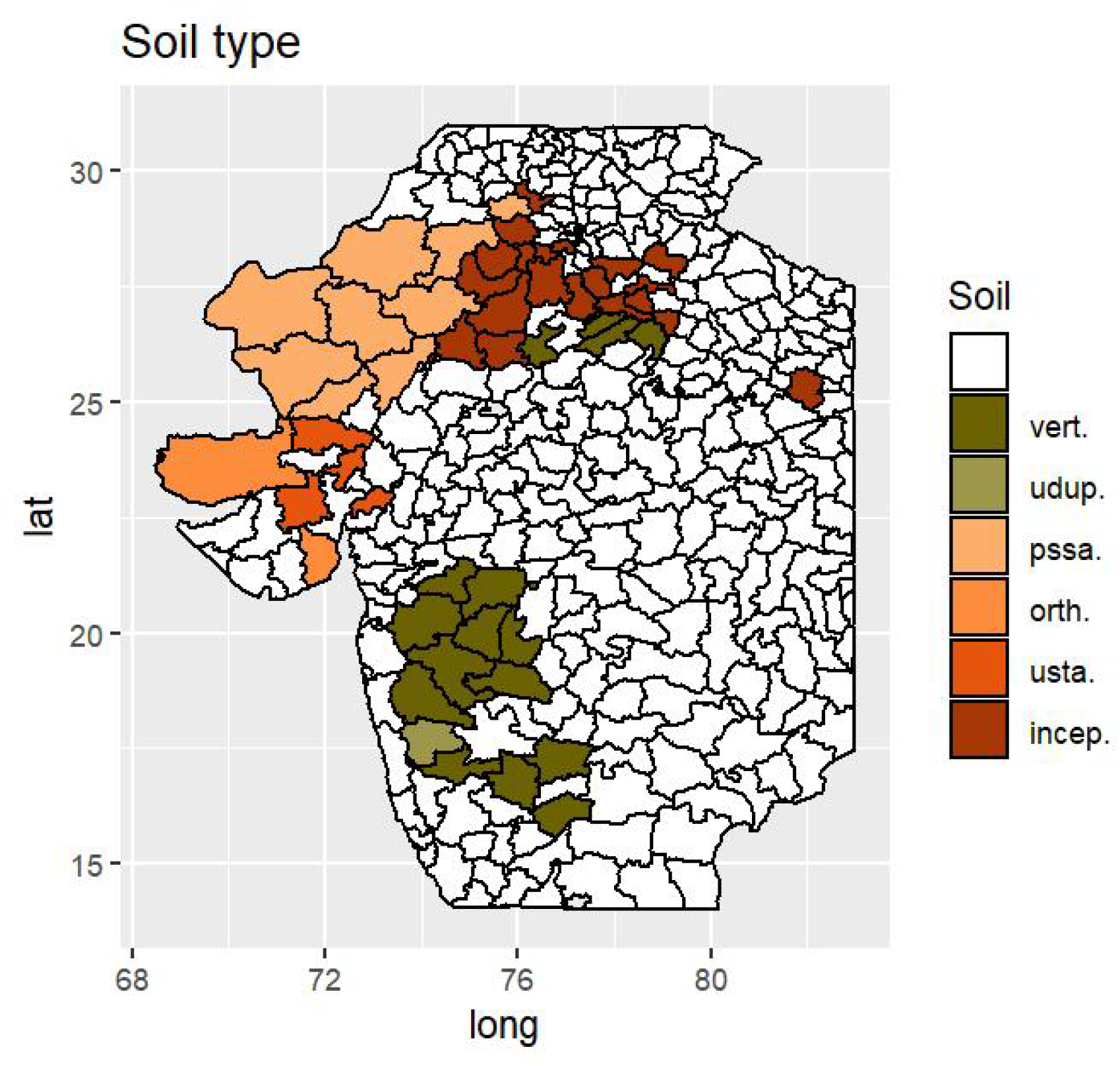
Majority soil type in the selected district representing 90 % of the total Kharif cultivated area between 1998-2017. Soil type: vert. = Vertisol; udup. = Udupt/Udalf; pssa. = Pssament; orth. = Orthid; usta. = Ustalf/Ustoll; incep. = Inceptisol

## S2: Crop model optimized parameter inference (OPI) illustratiom

**Figure 2:**
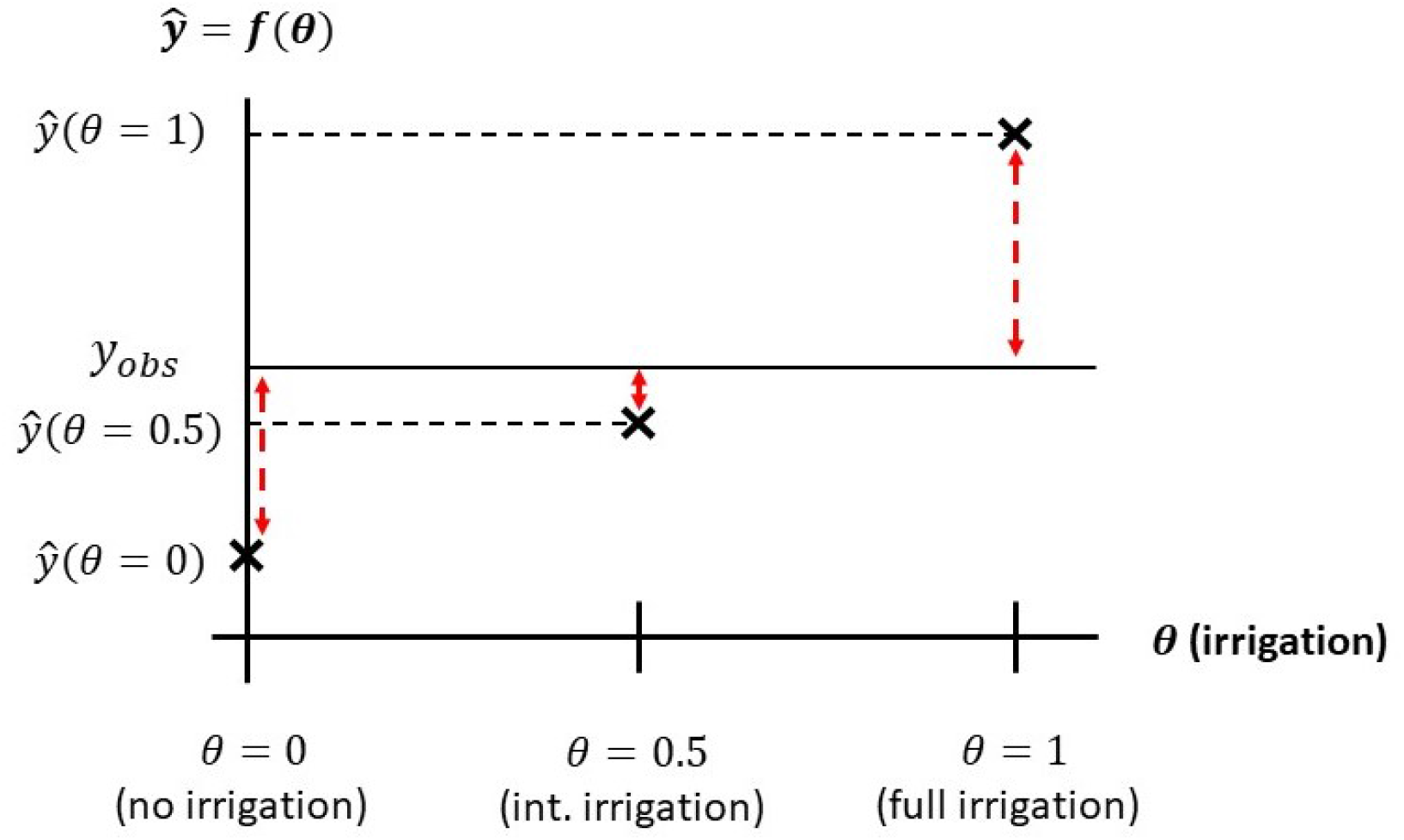
Illustration of the parameter inference strategy by comparing the CM output (*ŷ*) obtained for different values of a parameter (*θ*, here irrigation) with the observed or true value *y*_*obs*_. Here, we can notice that the parameter value (*θ* = 0.5) produces the output with the lowest difference compared to the true value. Therefore, assuming that an intermediate level of irrigation seems to be the most probable scenario

## S3: Reference selection of crop model parameter followimg the ‘imformed strategy’

**Table 1:** Crop model parameters

| Parameter | A1 zone | A zone | B zone |
| --- | --- | --- | --- |
| Model | New | New | New |
| Soil water content [mm/cm] | 0.6 (sandy) | 0.9 (clay/loam) | 1.3 (vertisol) |
| Soil depth [cm] | 120 (medium) | 120 (medium) | (medium) |
| Sowing date | 1-15 July (av.) | 1-15 July (av.) | 1-15 July (av.) |
| Sowing density [pl./m<sup>2</sup>] | 12 | 18 | 18 |
| Genotype (variety) | HHB67-2 | 9444 | 9444 |
| Fertilisation [kg N/ha] | DLD data* | DLD data | DLD data |
| Irrigation | DLD data | DLD data | DLD data |

\*: For fertilisation and irrigation, we set the parameter value to the one that was the closest to the average level of fertilisation and irrigation as observed in the DLD data over the considered period (1998-2017).

## S4: B zone area share, area trend and production compared to other crop

**Figure 3:**
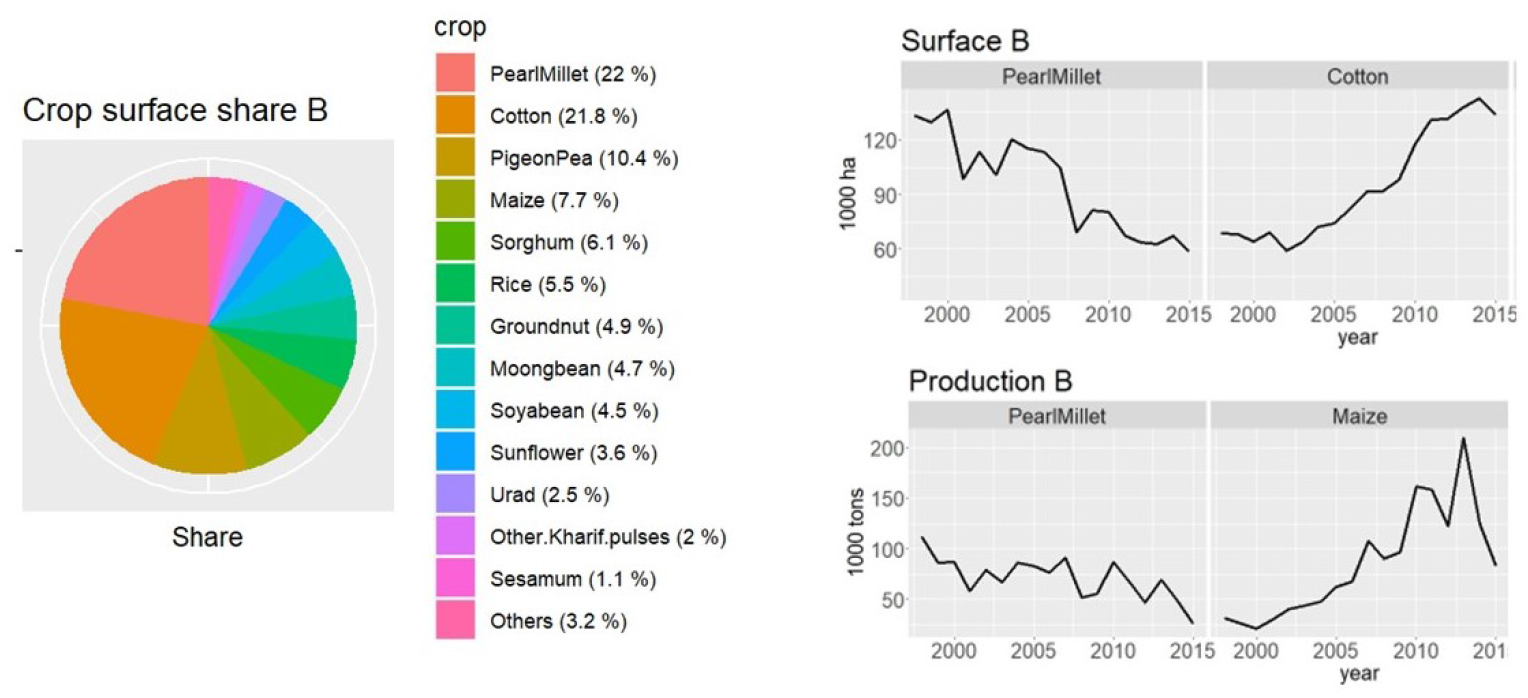
**Left part:** crop area share of pearl millet and other main crops cultivated in the B zone (1998-2015). **Right upper part:** surface trend of pearl millet and cotton (1998-2015). **Right lower part:** comparison of the pearl millet production trend compared to maize (1998-2015).

## S5: G zone area and production trend compared to other crops

**Figure 4:**
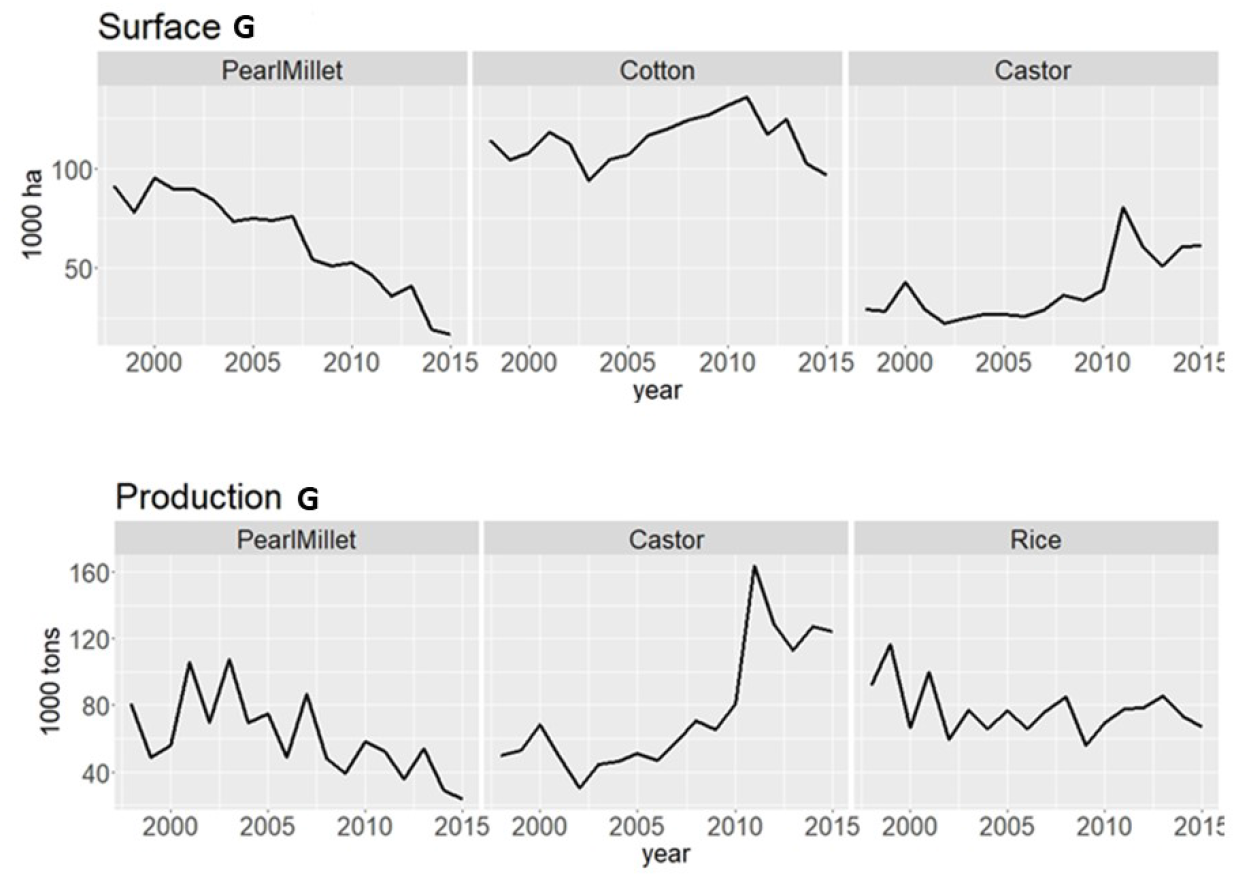
**Upper panel:** area trend of pearl millet, cotton and castor (1998-2015). **Lower panel:** comparison of the pearl millet production trend compared to castor and rice (1998-2015)

## S6: Influence of the crop model parameters on the crop model predictiom ability

We calculated the effect of the crop model parameter on the correlation (crop model prediction ability) by doing a linear regression of the different parameter options on the correlation between the observed and predicted yield. Those results allow the assessment of which parameter has an influence on the prediction ability, so which parameter should be correctly specified to model the yield trend with a good correlation.

From this analysis, we could notice that the version of the model used had a significant impact on the prediction ability in A1 and AE1 zones. In those zones, the new model performed significantly better. The correct specification of irrigation was also very influential in A1 and AE1. Simulation with intermediate and full irrigation gave less goodness of fit. The soil water content (PAWC) was also an influential parameter, especially in AE1, B, and G. Finally, the variety specification was also influential in AE2, B and G, where the use of improved variety (HHB67 or 9444) seems to be an important parameter for good prediction ability.

**Table 2:** Crop model parameter influence on model goodness of fit (correlation) in the new TPE as the estimated linear regression coefficient (B) and the R squared statistics over all simulations

|  | A1 |  | AE1 |  | AE2 |  | B |  | G |  |
| --- | --- | --- | --- | --- | --- | --- | --- | --- | --- | --- |
| var. | B | $R^2$ | B | $R^2$ | B | $R^2$ | B | $R^2$ | B | $R^2$ |
| Model:new | 0.00 | 0.14 | 0.00 | 0.12 | 0.00 | 0.03 | 0.00 | 0.08 | 0.00 | 0.06 |
| Model:old | -0.18 |  | -0.19 |  | 0.02 |  | 0.04 |  | -0.07 |  |
| PAWC:1.3 (clay) | 0.00 | 0.03 | 0.00 | 0.07 | 0.00 | 0.06 | 0.00 | 0.09 | 0.00 | 0.06 |
| PAWC:0.9 (loam) | 0.02 |  | 0.12 |  | 0.07 |  | 0.11 |  | 0.11 |  |
| PAWC:0.6 (sand) | 0.04 |  | 0.14 |  | 0.06 |  | 0.12 |  | 0.11 |  |
| SoilDepth:180 | 0.00 | 0.00 | 0.00 | 0.00 | 0.00 | 0.00 | 0.00 | 0.00 | 0.00 | 0.00 |
| SoilDepth:120 | 0.00 |  | 0.00 |  | 0.00 |  | 0.00 |  | 0.00 |  |
| SoilDepth:60 | 0.01 |  | 0.03 |  | 0.00 |  | 0.00 |  | 0.00 |  |
| Sowing:early | 0.00 | 0.06 | 0.00 | 0.04 | 0.00 | 0.11 | 0.00 | 0.10 | 0.00 | 0.05 |
| Sowing:average | 0.01 |  | 0.00 |  | -0.07 |  | -0.05 |  | 0.03 |  |
| Sowing:late | -0.01 |  | -0.01 |  | -0.04 |  | -0.07 |  | 0.05 |  |
| Density:12 | 0.00 | 0.00 | 0.00 | 0.00 | 0.00 | 0.00 | 0.00 | 0.00 | 0.00 | 0.00 |
| Density:18 | 0.01 |  | 0.03 |  | 0.00 |  | 0.00 |  | 0.02 |  |
| Density:24 | 0.02 |  | 0.04 |  | 0.00 |  | 0.00 |  | 0.03 |  |
| Variety:Landrace | 0.00 | 0.02 | 0.00 | 0.03 | 0.00 | 0.07 | 0.00 | 0.08 | 0.00 | 0.09 |
| Variety:HHB67 | -0.02 |  | 0.02 |  | 0.12 |  | 0.11 |  | 0.09 |  |
| Variety:9444 | 0.01 |  | 0.07 |  | 0.09 |  | 0.13 |  | 0.15 |  |
| Fertilizer:0 | 0.00 | 0.00 | 0.00 | 0.00 | 0.00 | 0.02 | 0.00 | 0.01 | 0.00 | 0.01 |
| Fertilizer:60 | 0.00 |  | 0.02 |  | 0.03 |  | 0.03 |  | 0.03 |  |
| Fertilizer:100 | 0.01 |  | 0.04 |  | 0.07 |  | 0.06 |  | 0.06 |  |
| Irrigation:off | 0.00 | 0.17 | 0.00 | 0.15 | 0.00 | 0.02 | 0.00 | 0.04 | 0.00 | 0.06 |
| Irrigation:int | -0.06 |  | -0.06 |  | -0.01 |  | 0.00 |  | -0.01 |  |
| Irrigation:on | -0.20 |  | -0.24 |  | -0.05 |  | -0.08 |  | -0.12 |  |

## S7: Comparison between observed data and parameters values determined with the OPI strategy soil parameters

**Figure 5:**
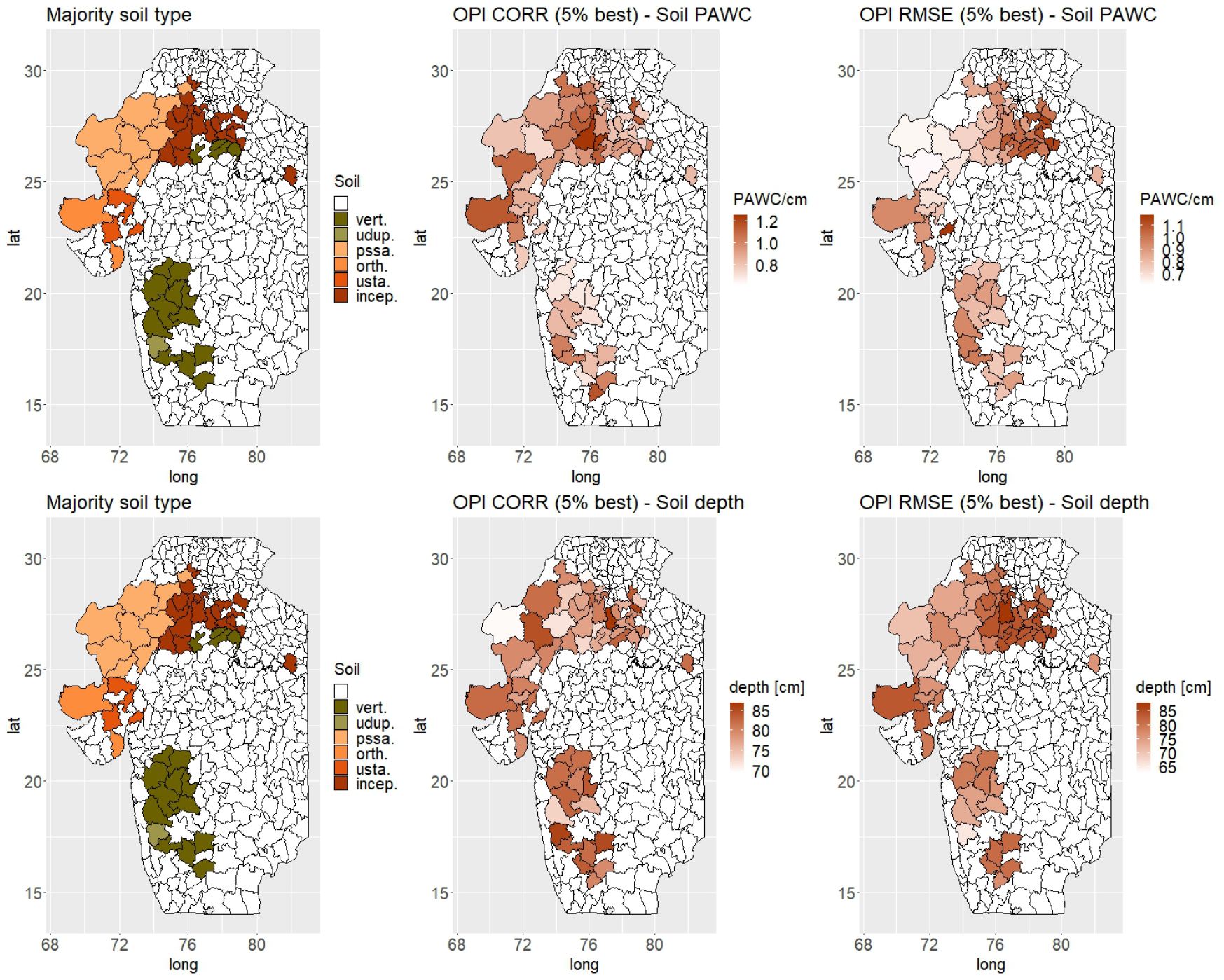
Comparison of the OPI parameter values for soil PAWC (**upper panel**) and depth with the type of soil (**lower panel**). The OPI parameters were the weighted average of the parameters values from the set of parameter with the largest predictive ability according to correlation (5% best results). We also included the results about parameter inferred using the root means square error and the same criteria (average parameter value of the 5% combination with the lowest RMSE). Using the pattern obtained with the RMSE was more consistent with our expectation of soil water content and depth in the observed majority soil types than the one obtained with correlation

## S8: Comparison between observed data and parameters values determined with the OPI strategy management parameters

**Upper panel:** comparison of the OPI parameter values for variety with DLD observation about the surface of pearl millet under improved variety. We converted the selected variety with the OPI method using the following scores: landrace (0), HHB67-2 (0.3), 9444 (1). **Middle panel:** comparison of the OPI parameter values for sowing date with the Number of days starting from 1 June to reach 25 mm of rain. **Lower panel:** Comparison of the OPI parameter values for plant density with the average cumulated rain during kharif.

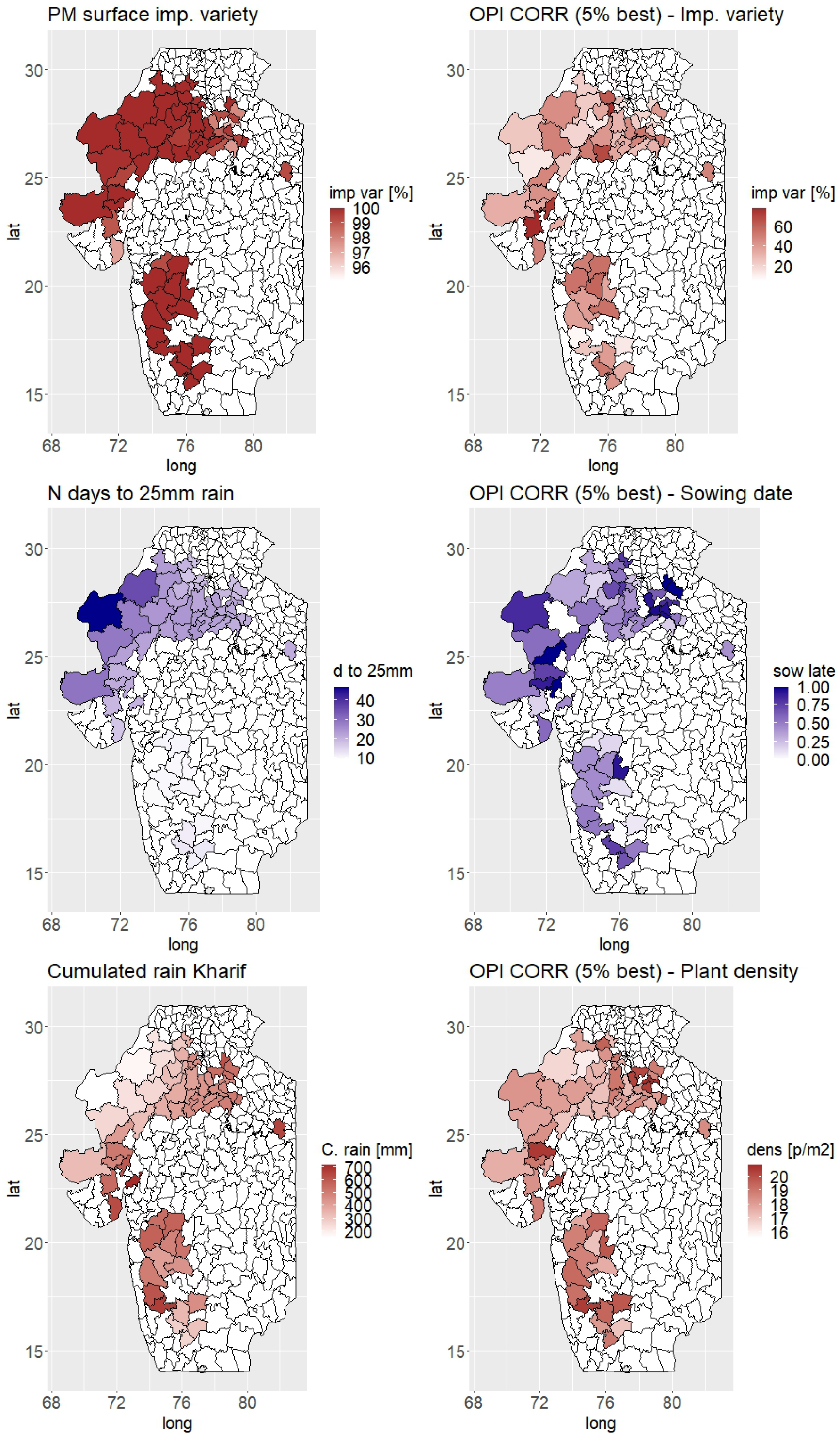

## S9: Pearl millet area trend over seasons in the G zone

**Figure 6:**
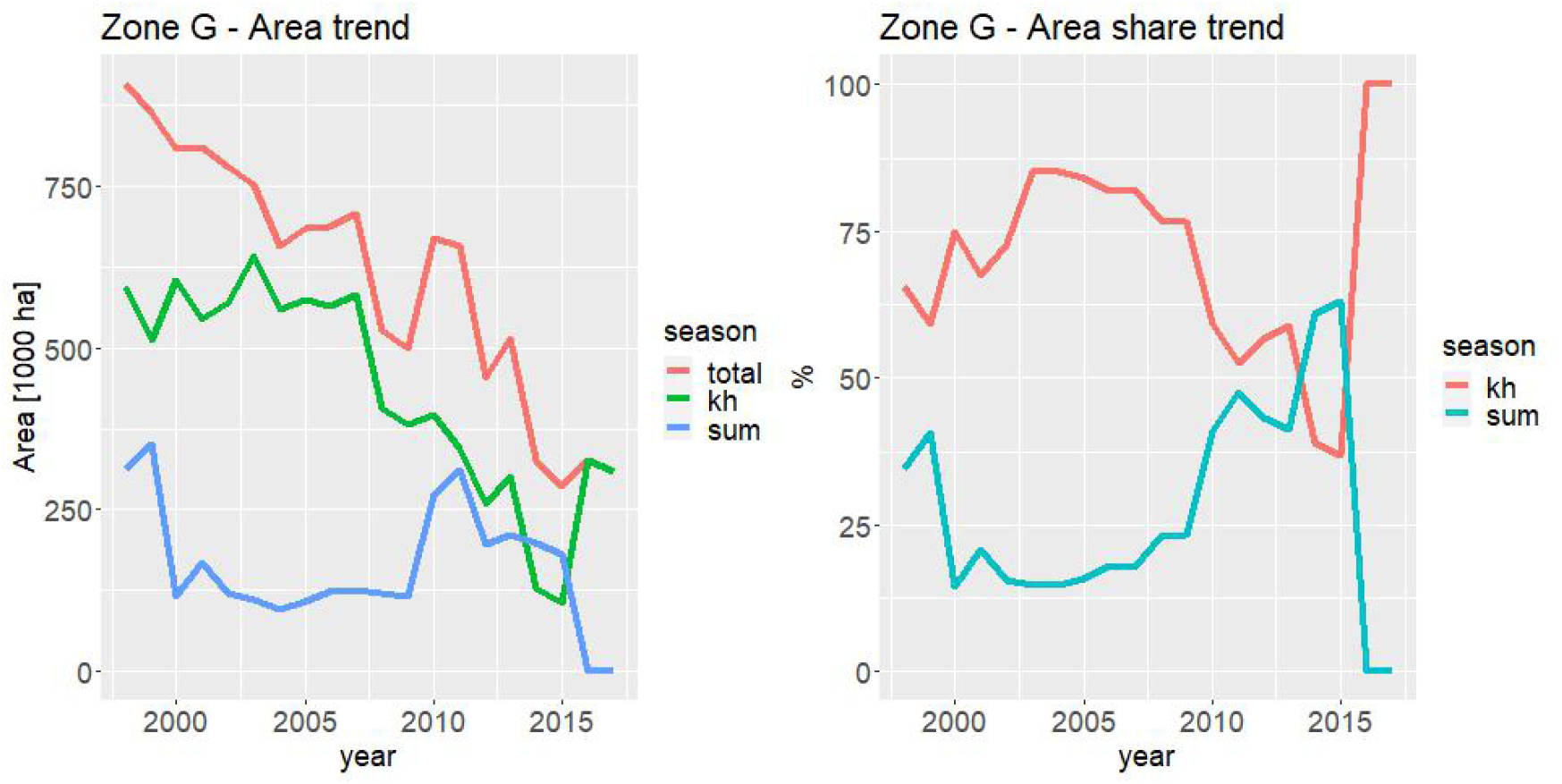
**Left panel:** Absolute value pearl millet cultivated area in the G zone during the kharif, summer and overal (kharif + summer) seasons. **Right panel:** Relative proportion of pearl millet cultivated during the kharif and summer season. We can notice that starting from around 2000, the proportion of area cultivated during summer increase to reach more than 50 %. The sudden drop at the end of the time series is potentially due to incomplete information about the summer season in the last years.

## S10: Pearl millet area trend over seasons in theAE2 zome

**Figure 7:**
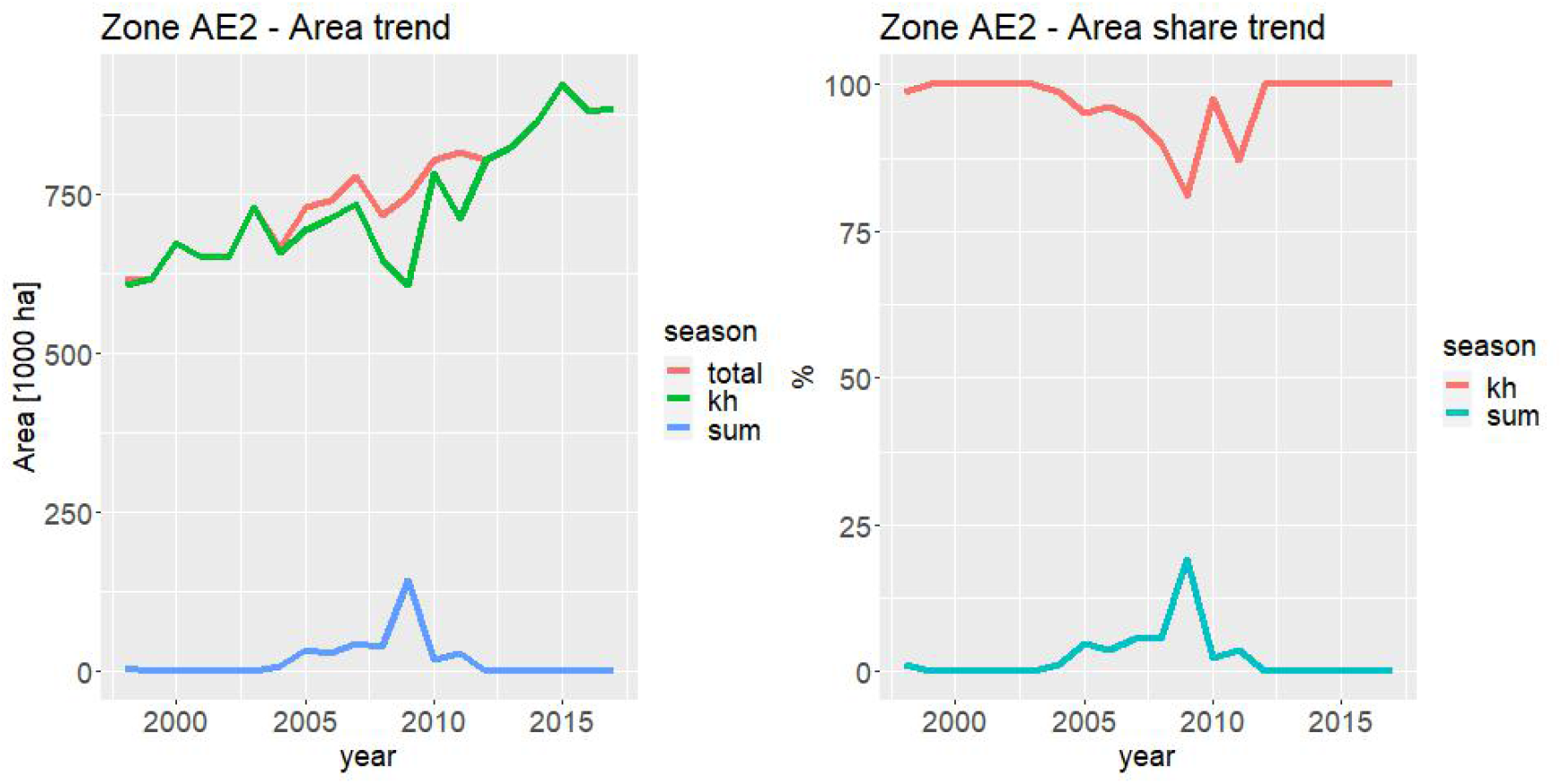
**Left panel:** Absolute value pearl millet cultivated area in the AE2 zone during the kharif, summer and overal (kharif + summer) seasons. **Right panel:** Relative proportion of pearl millet cultivated during the kharif and summer season.

